# Why link diverse citizen science surveys? Widespread arboreal habits of a terrestrial amphibian revealed by mammalian tree surveys in Britain

**DOI:** 10.1101/2022.02.27.482211

**Authors:** Silviu O. Petrovan, Nida Al-Fulaij, Alec Christie, Henry Andrews

## Abstract

Terrestrial anurans, with their typically short limbs, heavy-set bodies and absent claws or toe pads are incongruous tree climbers, but even occasional arboreal locomotion could offer substantial advantages for evading predators or accessing new shelter or food resources. Despite recent interest, arboreal behaviour remains rarely and unsystematically described for terrestrial amphibians in Europe, likely due to fundamental differences in survey methods and therefore a lack of field data. However, other taxa surveys specifically target trees and tree cavities. We undertook collaborations and large-scale data searches with citizen science projects surveying for arboreal mammals in Britain to investigate potential tree climbing by amphibians at a national scale. Remarkably, we found widespread arboreal usage by amphibians in England and Wales, with occupancy of hazel dormouse (*Muscardinus avellenarius*) nest boxes, tree cavities investigated as potential bat roosts and even a bird nest by common toads (*Bufo bufo*), but few additional records of frogs or newts. Toads are potentially attracted to tree cavities and arboreal nests because they provide safe and damp microenvironments which can support an abundance of invertebrate prey but the importance of such tree microhabitats for toad conservation remains unknown. Possible interactions with arboreal mammals are also unclear, but such mammals and even some birds may benefit from the occasional presence of toads if they feed on the mites and other arthropods that frequently infest their nests. We encourage expanding and linking of unrelated monitoring surveys and citizen science initiatives as valuable tools for investigating ecological traits and interactions.

## Introduction

Arboreal amphibians are vastly better represented in tropical compared to temperate regions, with the maximum diversity they reach in the tropics linked to the patterns of high precipitation and low annual seasonality as well as the variations in vegetation structure and the more complex habitat microniches in such environments (1). There is also substantial variation between temperate continental regions, with very few arboreal amphibian species in Europe compared with North America, and entirely represented in Europe by the genus *Hyla*, which was previously regarded as a single species - the European tree frog *Hyla arborea*.

Most arboreal amphibians use climbing as a way of locomotion in addition to jumping ability and have obvious morphological adaptations to facilitate surface climbing and movements on branches and leaves. Such adaptations in arboreal amphibians include toe disc modifications of the last phalanx to end with a hook as well as toe pads to allow gripping onto smooth surfaces but also longer digits, palm clasping, proportionally longer limbs and slender bodies and larger diapophyseal expansion, which allows greater fore-aft translation of the iliac shafts during climbing (2). Tree frogs also have strong abilities for attaching to a variety of surfaces using their versatile and complex toe pads which involves the secretion of mucus into the pad-substrate gap (3) but they likely rely on several attachment mechanisms given that they climb a wide diversity of natural surfaces and can vary greatly in size (4). By contrast, typical terrestrial amphibians are generally heavier, with squat bodies and proportionately shorter limbs (5; 6) and can produce substantially more eggs and prey on larger food items.

True toads, comprising the family *Bufonidae*, include both typical terrestrial hoppers but also riparian leapers (e.g. *Phrynoidis aspera*), terrestrial crawlers (e.g. *Melanophryniscus stelzneri*) and even several species of arboreal toads, in particular in SE Asia (e.g. *Pedostibes hosii*) (7). However, even some typical terrestrial toads which use hopping for locomotion, such as *Rhinella arenarum* from South America, were recently shown during climbing tests and morphological analyses of the limbs to be able to climb wooden structures of up to 90% inclination but were using different strategies compared with tree frogs, including flexing their fingers and toes to grasp at the substrate and displaying hooking and partial grasping (8). Climbing ability was also recently noted for other *Rhinella* toad species but based on few chance observations in the field (9; 10).

Temperate region toads are considered archetypal terrestrial amphibians and while there are some literature mentions of arboreal habits, these are typically rare observations of 1-2 individuals (11). These observations include the common toad *Bufo bufo* in Europe, one of the most widespread and common amphibians in Europe, which inhabits most of the continent, from the west coast of Britain to eastern Siberia and Kazakhstan (12). Species in other families of European terrestrial amphibians such as smooth newts *Lissotriton vulgaris* are also known to be able to climb vegetation and tree trunks, with some collated records from Denmark and Germany describing this behaviour (13). However, given that survey schemes for European amphibians focus almost exclusively on aquatic and ground–level terrestrial areas, there is an inherent inability to collect information on arboreal usage for this group, meaning our understanding remains very limited. There is substantial interest currently in managing European forests to benefit biodiversity including for providing and protecting microhabitats such as tree cavities for various groups such as bats, birds or insects (14; 15). Understanding if and how amphibians might use such arboreal microhabitats in trees could improve their conservation management and might be important for the broader implementation value of any such forestry focused management options.

Following a report about a toad in a dormouse nest box in England in 2016, and discussions with amphibian and mammal surveyors in Britain, it became apparent that while there are virtually no meaningful arboreal data for amphibians collected by herpetologists there might be broader potentially relevant survey data elsewhere. As an alternative to amphibian surveys, we used two major citizen science project schemes that focus on arboreal mammal monitoring in the UK to verify if such projects could contain valuable information on the arboreal occurrence of amphibians at a national level and, if so, to quantify and understand the extent, the potential reasons and implications for this behaviour at a national scale.

## Methods

To investigate potential tree climbing behaviour by amphibians we analysed data records from the main arboreal mammal survey projects in the UK, starting with the National Dormouse Monitoring Programme (NDMP). NDMP targets hazel dormouse, a nocturnal and rapidly declining arboreal species in the UK, which has become the focus of several conservation and reintroduction initiatives in England and Wales in the past decades (16). The scheme is supported and administered by the conservation NGO People’s Trust for Endangered Species (PTES) and between 1988-2014, 640 sites were monitored as part of NDMP, with a mean number of 77 boxes used to survey for dormice in sites with dormice presence (17), each checked a minimum of twice per year in spring and autumn, although some boxes are checked as often as monthly during the active season (April-November). Not all NDPM sites have dormouse populations and not all dormouse monitoring is part of this scheme, including those undertaken by ecological consultants in relation to planning proposals. NDMP guidance recommends that ideal monitoring sites should have 50 or more dormouse nest boxes, spaced 10-20 m apart in parallel lines, which should also be 10-20 m apart (18; 19). Dormouse nest boxes are wooden boxes similar to bird nest boxes but with the 3.5 cm entrance hole positioned immediately near the supporting tree or branch, and should be placed ideally 120-150 cm off the ground on coppiced hazel trees (*Corylus avellana*) where possible, or other shrubs or young trees well linked to the adjacent understorey and canopy (18). As dormice are legally protected monitoring requires a licence. The standardised NDMP recording form asks information on other mammals present in the nest box (e.g. mice or voles) but not specifically for other animal species (19).

Given that it was assumed that amphibian records were not consistently reported in the standardised forms by dormouse surveyors, a short online data request was sent to the NDMP surveyors by PTES in September 2016 and again in May-June 2021, asking for information about any amphibian records noted by observers during the nest box monitoring scheme. Data submissions were checked and some surveyors were additionally contacted to verify site details or to ask additional information. Some dormouse surveyors sent records from monitoring outside of NDMP, using both nest boxes and hair tubes.

Secondly, we investigated datasets collected for other arboreal mammals in the UK and in particular, bats. The citizen science initiative behind the Bat Tree Habitat Key (BTHK) project offers a publicly available but site anonymised database (20). The project began officially in 2010 and the majority of records were made in the period 2015-2019 but its associated database has records dating back to 2002. Surveyors, including trained and licensed volunteers and professionals, identify and surveys trees across the UK, searching for potential roost features (PRFs), and describe them in detail using standardised forms to record physical characteristics and environmental information. Records span all months and some PRFs have been subject to monthly inspections over three years. However, as with other citizen science projects, the BTHK project is mostly supported by qualified people recording data in their local woodlands and in their own free time. As a result, records tend to be biased toward periods when bats were present. Tree and site selection criteria vary, with some structured surveys of a particular area of wooded habitat where surveyors tried to map all the PRFs for repeat inspection as part of a Bat Group project, while others have radio-tracked bats to their roosts and catalogued the PRF as part of a research project, or have recorded roosts during ecological consultancy surveys (although these records are in the minority), and some volunteers just take their endoscope when walking in local woodlands to search and record PRFs they can access from the ground as they come across them. Collated data in the standardised BTHK forms include survey dates, tree location, tree species, tree height and DBH (diameter at breast height), PRF entrance height, the internal dimensions and the environment offered (e.g., apparent humidity, substrate texture and even smell) as well as any species using it; primarily bats but also any other mammals (e.g. squirrels), birds, arthropods, gastropods, other species and signs of animal usage such as bird or mammal nests. As bats are also protected species BTHK surveys operate under specific bat licences.

Most tree cavity inspections can be performed from the ground, but some bat surveyors are qualified to access PRFs in the canopy using specialist equipment, such as ropes or mechanical elevating work platforms. The PRFs are investigated using camera endoscopes (such as the Ridgid CA series or NHBS Explorer Premium). The endoscope lenses have integral LED lamps and live view is visible to the surveyor on a screen, so disturbance is controlled. In addition, the units allow the surveyor to record video footage and photographs of the inspection for later data verification and storage. This means that the numbers of bats and their species can be verified later, thus minimising the duration of the inspection. It also allows advice to be sought for other species that require specialist knowledge, such as invertebrates.

Finally, we discussed our data collection project with other NGOs and groups of ecological consultants to verify the presence of additional records from pre-existing survey datasets.

### Statistical analysis

To investigate tree and tree cavity selection by toads in the BTHK dataset we compared trees and PRFs used by toads to those where toads were absent and used a Generalised Linear Mixed Model (GLMM) (21; 22) and a Gaussian/normal error family as we expect tree and PRF size measurements to follow this distribution. We used four separate models to investigate the variation in four response variables (tree height (m), DBH (cm), PRF height (m), and PRF entrance height (cm)) and whether this was explained by the explanatory binary variable of the presence of toads. All analyses were carried out in R (23) using the lme4 (24), multcomp (25), and MuMIn (26) packages. To compare tree size measurements (DBH and height) we aggregated the BTHK dataset by the unique identifier for each tree surveyed and used a random effect for the survey location (i.e., site name) to account for the fact that several trees were sampled within each survey location. For PRF size measurements (height and entrance height) we aggregated the BTHK dataset by the unique identifier of the PRF (a combination of the tree identifier and PRF number) and used two nested random effects, tree identifier within survey location, to account for the fact that sometimes several PRFs were sampled on the same tree, and several trees were sampled within each survey location. We determined p-values and modelling statistics by comparing the model with the term of interest (presence of toads) to a model without (i.e., an equivalent intercept-only model), and then conducting a likelihood ratio test.

We also compared the tree and PRF measurements for trees where toads were present to those in which slugs, snails, blue tits (*Cyanistes caerulaeus*), and woodlice (*Oniscidea*) occurred (as well as all other animals in the BTHK dataset labelled as ‘other). We selected these animal groups as they represented the four most recorded animal groups in the BTHK dataset (see Table 1). For all species data were recorded as presence absence in each PRF but for vertebrates the total number was recorded when there was more than one individual present. We used the same four model specifications as before (in terms of tree and PRF measurements and random effects) but used a different categorical explanatory variable that indicated the presence of toads, slugs, snails, blue tits, woodlice, or other animal groups. We used multiple comparison tests with the Tukey adjustment to test for differences in tree and PRF measurements between these animal groups. Plots of model residuals approximately followed a normal distribution and there were no strong patterns of residuals versus fitted model values, indicating modelling assumptions held. Marginal and conditional R^2^ values were computed for each model.

**Table 1.**
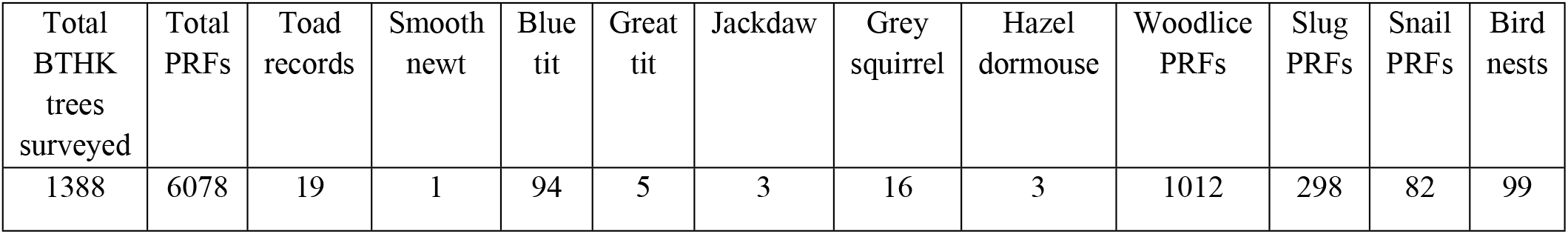
Total trees and tree cavities (PRFs) surveyed as part of the standardised monitoring in the Bat Tree Habitat Key scheme in the UK and comparisons of amphibian records with other animal species and bird nests identified.

| Total<br>BTHK<br>trees<br>surveyed | Total<br>PRFs | Toad<br>records | Smooth<br>newt | Blue<br>tit | Great<br>tit | Jackdaw | Grey<br>squirrel | Hazel<br>dormouse | Woodlice<br>PRFs | Slug<br>PRFs | Snail<br>PRFs | Bird<br>nests |
| --- | --- | --- | --- | --- | --- | --- | --- | --- | --- | --- | --- | --- |
| 1388 | 6078 | 19 | 1 | 94 | 5 | 3 | 16 | 3 | 1012 | 298 | 82 | 99 |

## Results

We identified and collated records of amphibians associated with dormouse surveys from 18 sites, with dates of observations spread between 2009 and 2019. Most records (30 individuals) came specifically from checking dormice nest boxes as part of NDMP, but one was from a recent but empty blackbird (*Turdus merula*) nest found in the tree while checking the dormouse nest box. Another record was from an ecological survey to verify the presence of dormice using hair tubes, with a toad using the hair tube, and one was from dormouse monitoring using nest boxes but not part of the national monitoring scheme. Although several amphibians were found in dormice nests inside nest boxes, none were observed simultaneously in the nest box or the tree cavity with arboreal mammals or birds in either of the survey schemes investigated. All amphibian observations from the dormouse survey scheme were linked to rural woodland areas located in England and Wales (Fig 1A).

**Figure 1.**
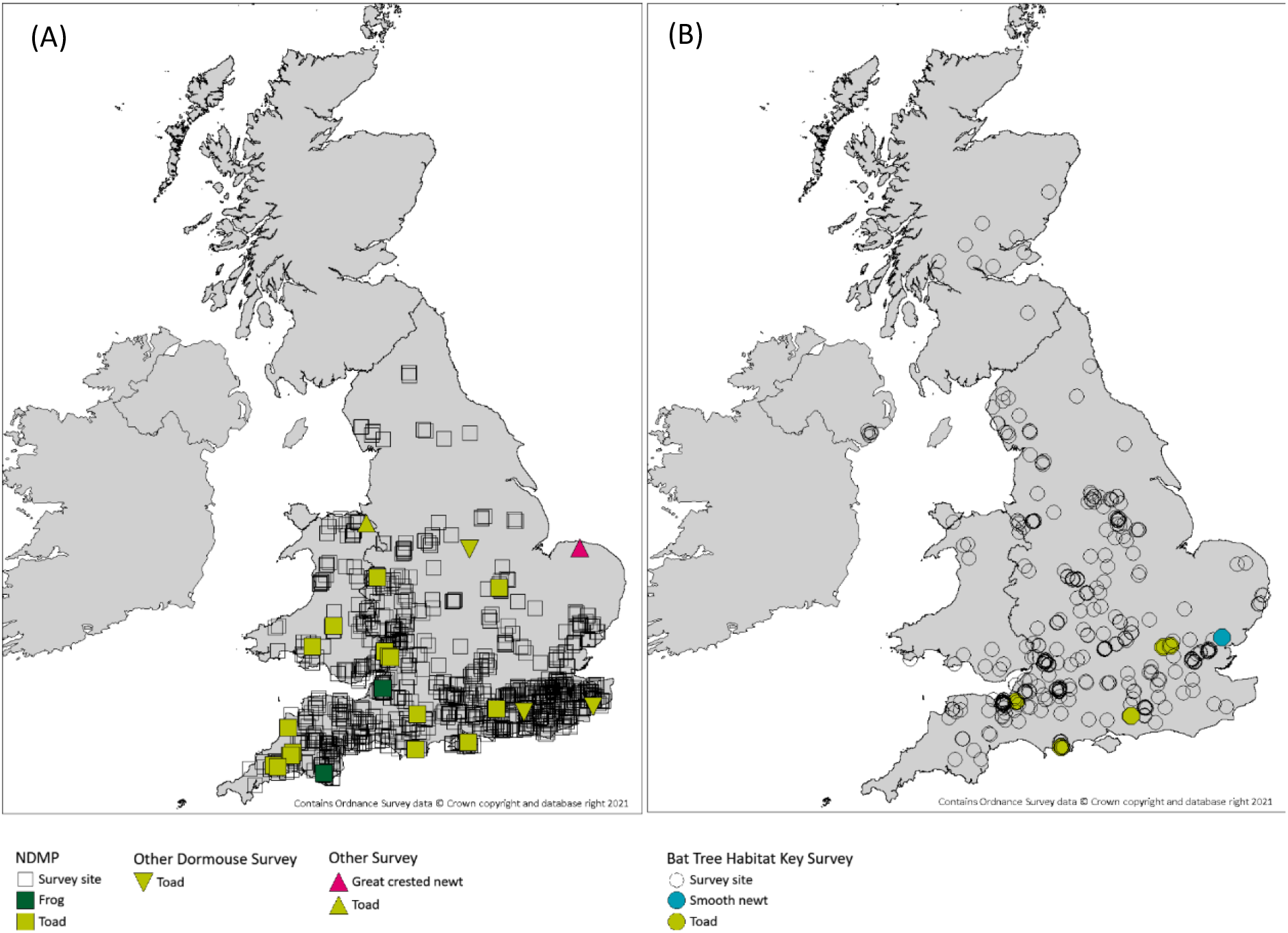
A. Survey sites and amphibian records as part of National Dormouse Monitoring project plus other single site arboreal mammal surveys. Note that in some cases there are multiple toad records in the same site. B. BTHK survey sites and amphibian records. Note that in some cases there are multiple toad records in different trees at the same site or in the same tree.

In addition, the 1,388 trees surveyed in the Bat Tree Habitat Key project generated a further 20 other amphibian records from 5 sites (Fig. 1B; Table 1), all from 2015-2019, including with multiple individuals. A distinct record came from a separate bat roost survey.

Most amphibians recorded were common toads but we also collected two records of common frog, *Rana temporaria* in dormouse nest boxes and two of newts, a smooth newt male and two great crested newts *Triturus cristatus*, found in tree cavities during bat surveys. One adult toad was discovered dead inside a dormouse nest box but the cause of death was unknown.

There was no obvious seasonal pattern in the distribution of amphibians in either next boxes or tree cavities, but of the total 54 amphibians recorded there were more observations in summer months May-July (54% of observations) compared to spring (March-May: 9%) or autumn (September-October: 37%).

Nest box height was sometimes not recorded in the NDMP database, as most sites include a substantial number of such boxes and the variation between them in terms of height is small as following guidance most are placed at “chest height” or between 120 to 150 cm height, to facilitate checking by volunteers. For the BTHK, where PRF height was recorded as standard, the mean height of cavities occupied by toads was 134 cm but there were records of 192 cm and 216 cm and the maximum recorded cavity height occupied by a toad was over 3 m, within a cavity with the entrance at 280 cm height in an oak tree and an additional 25 cm up above the entrance inside the feature (Figure 2A).

**Figure 2.**
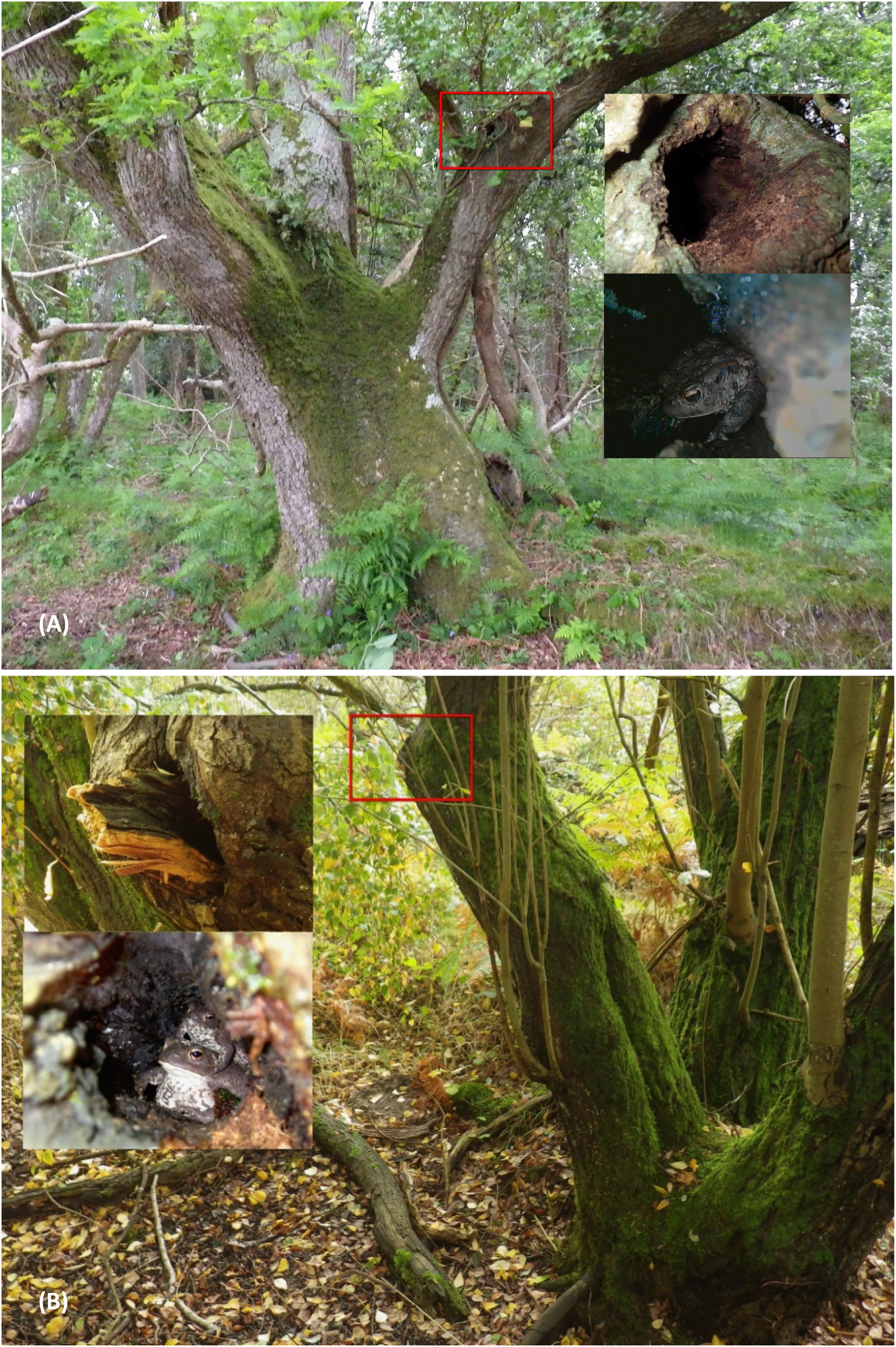
Total tree aspect, detail of PRF entrance and internal PRF image examples from BTHK. A the PRF (knot-hole type) is 2.8 m up the common oak and an adult toad, probably male, is visible in the endoscope image. B the PRF (tear-out type) is 0.93 m up the goat willow and two toads, an adult and a subadult are visible inside. Images Henry Andrews.

The average size of trees occupied by amphibians in BTHK (trees used by toads: average height 10.4 m, average DBH: 28.8 cm) was smaller compared with the wider dataset of surveyed trees (average tree height: 12.6 m, average DBH: 43.6 cm), with wide variation between groups of animals recorded in tree cavities (Figure 3).

**Figure 3.**
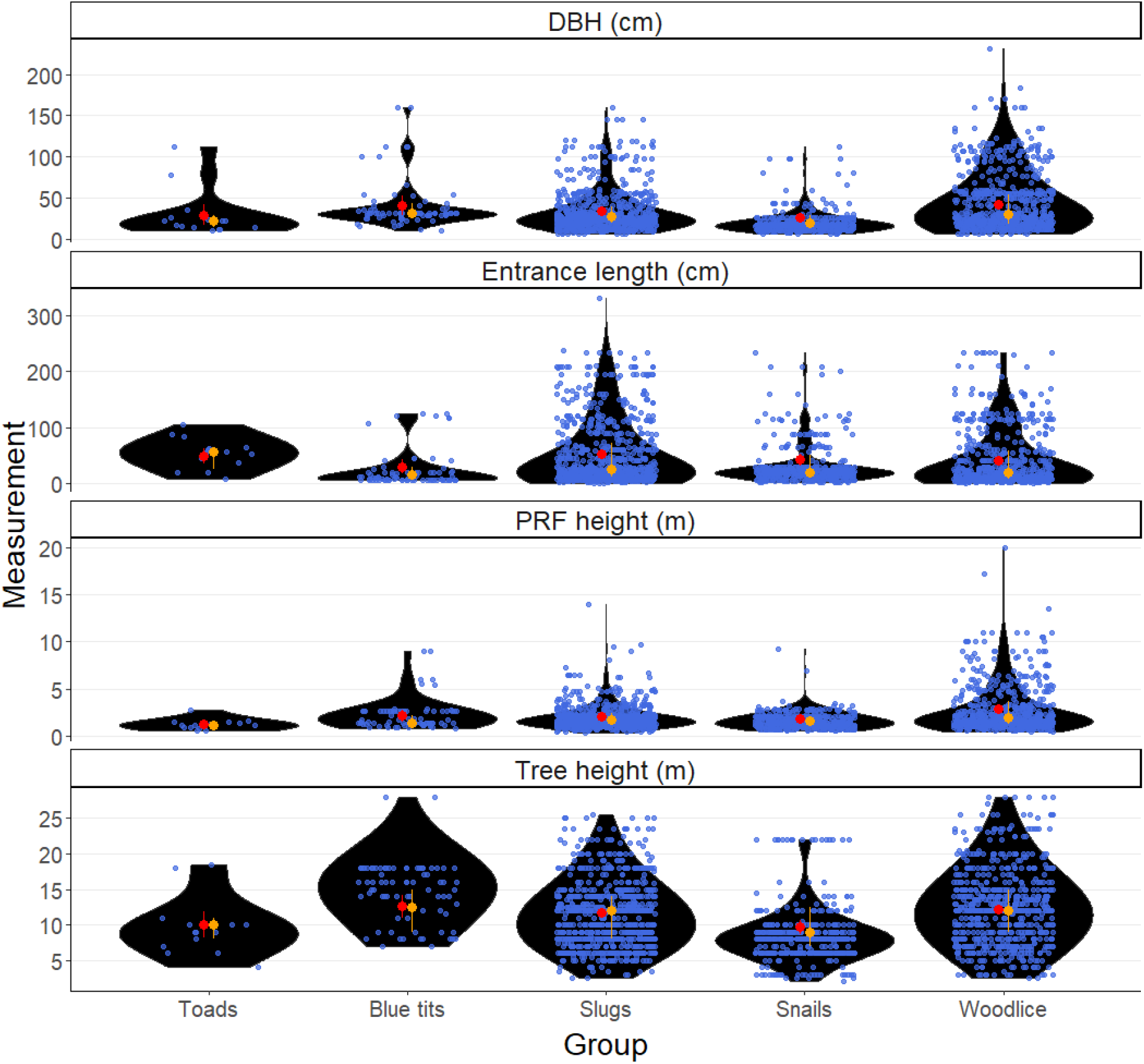
Comparative tree characteristics for multiple taxa recorded in tree cavities (PRFs) as part of the BTHK survey showing raw mean values plus 95% confidence intervals in red; median plus upper and lower quantiles in orange. To aid data visualisation, we have excluded four datapoints for PRF height greater than or equal to 13.5m (3 for woodlice and 1 for snails).

In the BTHK dataset, the number of toad records was small, which limited the statistical power of our models to detect differences in tree measurements between trees with toads versus other animals. The fixed effect of toad presence was poor at explaining the variation in different tree and PRF measurements (extremely low marginal R^2^ values, all less than 0.004) and it was clear that the random effects of survey location and tree identifier explained far more of the variation (higher values of conditional R^2^ values ranging from 0.30 – 0.88;). Nevertheless, summary statistics and high variability in tree and PRF measurements in the BTHK dataset supported our model’s inconclusive findings that toad selection of trees was similar compared to the wider dataset in terms of tree height (tstat: 0.71; pvalue: 0.98), DBH (tstat: 0.73; pvalue: 0.97), PRF entrance height (tstat: 0.97; pvalue: 0.93). Trees occupied by toads were also similar in height to those selected by blue tits, snails, slugs, and woodlice (Figure 3; Supplementary material S1), but there was an indication that snails were selecting lower height trees compared to the available trees (tstat: −2.86; pvalue: 0.04). There was no apparent pattern in the orientation of the entrances into PRFs used by amphibians, with three PRFs facing NW, three SE, three SW, four West, four East and one North.

Amphibians in BTHK were recorded in PRFs located in seven tree species: sycamore *Acer pseudoplatanus*, alder *Alnus glutinosa*, downy birch *Betula pubescens*, silver birch *Betula pendula*, hazel *Corylus avellana*, common oak *Quercus robur* and especially goat willow *Salix caprea*. Compared to the nearly 50 species surveyed overall in BTHK (including some hybrids and others identified only to genus level), the tree selection by amphibians was broadly similar to its availability in the dataset for some tree species, with of the two most common tree species surveyed in BTHK, sycamore and common oak, used by 17.6% and 11.8% of toads and represented 6.9% and 11% of all trees surveyed. However, there was a substantial difference apparent for goat willow, which was used by 35.3% of amphibian records despite representing only 1.1% of all trees surveyed in the BTHK project and suggesting positive selection for the environmental conditions associated with this tree species (e.g. damp or wet woodland). By contrast, pedunculate oak *Quercus petraea* was the third most frequently surveyed tree species in the project (33.6% of all surveyed trees) yet none of the PRFs surveyed for this tree species were used by amphibians. All trees used by amphibians were live trees.

## Discussion

Most animal species use a characteristic primary mode of locomotion for the majority of their daily activities, but several species were shown to be capable of expanding their locomotion mode in order to access atypical habitats or substrates, such as some European terrestrial rodents when climbing vegetation (27). Even if rarely used, this ability to adjust the movement type to access otherwise inaccessible areas may confer those individuals important or even critical advantages in particular situations such as during dispersal, when facing stressful environmental situations such as drought, fires or flooding, or during the generation of new ecological niches (8). The collated data from arboreal mammal surveys in Britain demonstrates that some amphibian species regularly climb trees in Britain and do so across their active period in the year, although with an apparent increase in summer and autumn months. While literature examples and discussions with experts indicated such behaviour and ability to climb vegetation for some newt species and especially the smooth newt (11), our collated dataset from nest boxes and tree cavities is overwhelmingly and unexpectedly comprised of common toad records.

Common toads are morphologically a typical terrestrial anuran, with short legs keeping the body close to the ground, slow walking or hopping movements and heavy body weight, especially for adult females, but which has been described as a “laborious climber” which can overcome many obstacles on its way (28), particularly during the spring migration to the breeding ponds (29). They are considered adaptable habitat generalists, inhabiting woodland, grassland, farmland and coastal areas, can tolerate some degree of urbanisation and often occupy artificial wetlands such as reservoirs or large man-made ponds (30) although it has suffered large scale declines in Britain in recent decades (31). Toads live overwhelmingly terrestrial lives, normally only found in water during breeding in March-April as adults and March-July as tadpoles, they hibernate on land, and usually spend daytime periods under dead wood or large rocks and emerging at night to ambush hunt woodlice, earthworms, slugs and ants. The preference for wooded habitat, in particular deciduous woodland, is well known for this species, with the probability of toad occurrence positively associated with the presence of nearby wooded habitat (32; 33). Yet, despite the fact that their biology and ecology are well documented and that it is universally described as a terrestrial species, there are rare instances documenting vegetation climbing in this species but they are either general and do not provide specific details (28) or refer to chance observations of 1-2 individuals (13). However, Gosá (34) recorded in northern Spain that local toads (now recognised as a separate species, *Bufo spinosus*) were using roots and low oak-trunk sections in an old-growth oak forest and collected over 200 observations of amphibians in 2000-2003 of such climbing behaviour, mostly *Bufo spinosus* (129 observations with an average climbing height of 39 cm and maximum height of 197 cm) but also *Alytes obstetricans* (66 observations at 34 cm average height, maximum height 135 cm) and *Rana temporaria* (9 observations, 14 cm average height, maximum 30 cm) and suggested this behaviour was linked to a search for humidity provided by moss growing on oak as records were rare during the wet season (March to early June) but increased during the dry period (September—October) (34).

While the 19 toad records in 1,388 trees surveyed (1.37% occupancy) and over 7000 tree cavity surveys in the BTHK database might suggest toads are relatively rare users of tree cavities, the numbers of toad records are comparable with the those for other vertebrate species in the same dataset (Table 1). For instance, several deciduous tree cavity nesting bird species with very large breeding populations in the UK such as blue tits, estimated at 3.6 million breeding territories, had only 94 records in BTHK. Even fewer records were collated for other common birds that tree cavities, including great tits (*Parus major*) with a UK breeding population estimated at 2.5 million pairs or jackdaws (*Corvus monedula*) with 1.4 million pairs (35). Only 99 additional BTHK records included empty bird nests in tree cavities. Altogether, the relatively small number of BTHK records of species known to often rely on tree cavities for breeding, such as blue tits, and their overall UK abundance numbering in the millions, plus the fact that there are 3.23 million hectares of woodland in the UK (36), containing perhaps 3 billion trees, suggest that the number of toads regularly using tree cavities in Britain could be substantial. As shown at the site with the highest numbers of observation (West Heath in Hampshire), the presence of suitable trees with tree cavities and large ponds nearby, might increase opportunities for tree habitat usage by toads. This matches well with the proposed conservation measures for common toads, that include increased density of both wooded and wet habitats (e.g. through pond and ditch creation) in farmland (32). That goat willow appeared particularly used compared to their low availability is not surprising given that this tree prefers wet areas, often bordering bodies of freshwater such as lakes. It is however important to note that that the overall sampling regime in our dataset was biased towards the survey of target species (i.e. hazel dormouse and bats) and as such these results are potentially not representative of the true habitat use of non-target species such as toads.

The spatial distribution of amphibians in dormouse nest boxes in our dataset is probably an artefact of the dormouse distribution area in Britain and the monitoring survey intensity for this species, which are mainly focusing on their remnant strongholds in southern England and southern Wales and the English-Welsh border (19). The Bat Tree Habitat Key tree monitoring database is more widely distributed in the UK, reflecting the broader distribution of tree-dwelling bat species in Britain compared with dormice (Fig 1B). However, a relatively similar distribution pattern was apparent for the amphibian records collated as part of BTHK project, despite the wider, if unequal spatial coverage across Britain and even in Northern Ireland.

Anurans can use a range of movements on the ground, including leaping, walking, crawling or hopping. However, while arboreal species have no difficulty to switch to terrestrial locomotion (as most anurans are capable of hopping), the reverse is far more problematic for terrestrial anurans with short limbs and a heavy body, and thus cases of terrestrial frogs or toads climbing trees or cliffs remain rare (8). What is particularly remarkable in our dataset is the height of several observations, with a record of a toad in a tree feature with the entrance at 2.8 m height from the ground. By comparison, of the four individuals of *Rhinella margaritifera* and one individual of *R. castaneotica* recorded above ground level on vegetation, one was at 130 cm above the ground while the others were at 32 cm, 45, 75 and 102 cm above ground level (9). For another terrestrial anuran capable of climbing, the catastrophically invasive cane toad *R. marina* in Australia, Hudson et al. (37) found strong differences in climbing ability associated with sex and relative limb length, but also population of origin, with longer-limbed male individuals as better climbers within each population. Yet, the climbing ability of cane toads appeared primarily driven by the local environmental conditions that supported or rewarded such arboreal activity (37).

Few European terrestrial amphibians are known to climb tree trunks and low branches but smooth newts have been recognised as capable climbers (13). The reasons why they do so remain unknown and the extent of this behaviour might be underestimated by our data which did not include surveys of shrubs. While European newts have lungs, they are superficially similar to plethodontid salamanders (found mostly in temperate and tropical Americas) which are known to have substantial arboreality, with some 45% of all non-aquatic species being either arboreal or facultative arboreal (38). Yet, even for plethodontid salamanders the prevalence of arboreal behaviour remains insufficiently recognized and often reliant on opportunistic observations (38), thus hampering adequate links with species ecology and conservation management of their environments.

Our systematic field surveys of dormouse nest boxes and unsystematic but large-scale surveys of tree cavities demonstrate that common toads, although apparently poorly suited morphologically to this locomotion type, are in fact capable of extensive tree climbing. Common toads presumably achieve this by using the fingers and toes to perform sufficient substrate gripping to allow them to climb arboreal environments, both for relatively flat and steep angle large tree trunks as well as near-vertical small diameter tree trunks. However, why apparently substantial numbers of adult toads climb trees, how long they remain there, and how they select trees with cavities or arboreal nests remains unknown. An arboreal niche might allow toads opportunities to survive either as a resting site where predators can be avoided, or as novel foraging areas (8; 39) compared to the ground level where they risk being hunted or parasitized. The toadfly *Lucillia bufonivora* is the obligate agent of myiasis in amphibians and an important specific parasite of common toads, found in both open habitats and shaded woodland in different European studies (40). In Britain, most toadfly records are from England (41), yet even there it is considered rare, perhaps a consequence of the recent declines of its main hosts, the common toad. Similarly, barred grass snakes (*Natrix helvetica*) are the main predator of toads and are common and widely distributed in England and Wales. They possess the ability to consume common toads as tadpoles and adults, despite toads being poisonous to other species. Both toadfly and grass snakes are largely absent in Scotland, where we also did not record any observations of amphibians in tree cavities. However, this could also be explained by the biases in the datasets we analysed. The hypothesis that toads climb trees more often in areas with high predator or parasite risk remains untested but could be a topic for future studies.

Our findings on the climbing ability of toads also have practical conservation relevance since toads often fall into road drains and gully pots. A central mitigation solution is to install perforated metal or mesh “ladders” to allow escape from such traps (42) and a good climbing ability is therefore crucial.

We can only speculate as to the exact reasons for the presence of toads in dormice nests boxes. However, some information does exist on amphibians using arboreal mammal nests, such as arboreal salamanders *Aneides lugubris* and *A. ferreus* utilizing *Arborimus spp*. vole nests up to at least 20 m high in forest canopy in western USA (43). In six of the ten cases, both salamander species and voles were present at the same time and authors suggested that the presence of salamanders may benefit the voles by feeding on the mites and dipterans which may parasites the voles (43).

Currently, survey limitations and lack of appropriate field data are hampering our ability to investigate the arboreal ecology and behaviour of common toads. For example, much like other British amphibians, the nocturnal and generalist nature of common toads means that nearly all surveys of this species are undertaken during the breeding time, when adults congregate at aquatic sites. Substantially less survey effort is targeted at their terrestrial habitats given that toads can inhabit many different habitats and are difficult to detect (32). This is a common problem for amphibian surveys but also for other nocturnal species surveys, where observations are generally biased towards sites or times that facilitate observations. However, as shown in this study, there is untapped potential to use data from surveys targeted at particular species, such as from volunteer-led surveys and citizen science, to answer interesting questions for other, non-target species. For example, in the UK, where citizen science has a long history and diversity of projects (44), one of the largest structured datasets of mammal records comes from the Breeding Bird Surveys (BBS). The BBS is run with volunteers and coordinated by the British Trust for Ornithology, generating important understanding of mammal distribution and abundance trends, although there are some careful considerations to consider during data verification and expert validation of spatial outputs (45).

Arboreal locomotion and occupancy of tree cavities and nests in European forests by terrestrial amphibians such as common toad appears a much more common phenomenon than previously thought, yet this apparently widespread behaviour remains largely unrecognised and the drivers behind it are unknown. The fact that standardised survey data has existed unused for nearly a decade in Britain from separate monitoring projects should act as an incentive for other researchers to investigate such collaborations. Future citizen science should look beyond distribution and abundance data and target complex species interactions (46); collecting and integrating diverse citizen science datasets across taxa groups could provide valuable datasets for further study.

## Acknowledgements

We are grateful to the countless volunteers that have contributed data to these arboreal mammal projects; their efforts have made this analysis and data collation possible. In addition, we want to thank Roger Downie, Arnold Cooke and Robert Oldham for useful discussions and advice from their decades of studying frogs and toads in the field and their comments on earlier drafts of this work.

## Ethical Statements

Funding: the authors declare that no funds, grants, or other support were received during the preparation of this manuscript. Conflict of interest: none. Ethical approval and informed consent: All observations followed protocols for protected species and the ethical standards considered for these licenses. No animal testing or experimentation took place.

## Data availability

All BTHK data is openly available http://battreehabitatkey.co.uk/?page_id=18 but site location is excluded. Overall site location data is not provided for any observations given the protected status of the included surveyed species (bats and dormice) and the best practice for records of such species.

---

**Table S1 (Supplementary material).** Log likelihood test for a general linear mixed model comparing a model with the term of interest (i.e., toads present), with model without the term of interest.

| Measure type | Model | npar | AIC | BIC | logLik | Deviance | ChiSq | Df | Pr(>Chisq) |
| --- | --- | --- | --- | --- | --- | --- | --- | --- | --- |
| Tree height (m) | Model without toad variable | 3 | 8218.343389 | 8234.032905 | -4106.171695 | 8212.343 | NA | NA | NA |
| Tree height (m) | Model with toad variable | 4 | 8220.187905 | 8241.107261 | -4106.093953 | 8212.188 | 0.155484 | 1 | 0.693349 |
| DBH (cm) | Model without toad variable | 3 | 13022.92319 | 13038.59964 | -6508.461597 | 13016.92 | NA | NA | NA |
| DBH (cm) | Model with toad variable | 4 | 13024.65678 | 13045.5587 | -6508.328389 | 13016.66 | 0.266416 | 1 | 0.605746 |
| PRF height (m) | Model without toad variable | 4 | 7294.0685 | 7315.619337 | -3643.03425 | 7286.069 | NA | NA | NA |
| PRF height (m) | Model with toad variable | 5 | 7294.172352 | 7321.110898 | -3642.086176 | 7284.172 | 1.896149 | 1 | 0.16851 |
| Entrance height (cm) | Model without toad variable | 4 | 17442.09283 | 17463.53325 | -8717.046416 | 17434.09 | NA | NA | NA |
| Entrance height (cm) | Model with toad variable | 5 | 17444.08187 | 17470.88239 | -8717.040935 | 17434.08 | 0.010962 | 1 | 0.916613 |

## Notes

### Competing Interest Statement

The authors have declared no competing interest.

